# Data-driven and interpretable machine-learning modeling to explore the fine-scale environmental determinants of malaria vectors biting rates in rural Burkina Faso

**DOI:** 10.1101/2021.04.13.439583

**Authors:** Paul Taconet, Angélique Porciani, Dieudonné Diloma Soma, Karine Mouline, Frédéric Simard, Alphonsine Amanan Koffi, Cedric Pennetier, Roch Kounbobr Dabiré, Morgan Mangeas, Nicolas Moiroux

## Abstract

**Background:** Improving the knowledge and understanding of the environmental determinants of malaria vectors abundances at fine spatiotemporal scales is essential to design locally tailored vector control intervention. This work aimed at exploring the environmental tenets of human-biting activity in the main malaria vectors (*Anopheles gambiae s.s.*, *Anopheles coluzzi*i and *Anopheles funestus)* in the health district of Diébougou, rural Burkina Faso.

**Methods:** *Anopheles* human-biting activity was monitored in 27 villages during 15 months (in 2017-2018), and environmental variables (meteorological and landscape) were extracted from high resolution satellite imagery. A two-step data-driven modeling study was then carried-out. Correlation coefficients between the biting rates of each vector species and the environmental variables taken at various temporal lags and spatial distances from the biting events were first calculated. Then, multivariate machine-learning models were generated and interpreted to i) pinpoint primary and secondary environmental drivers of variation in the biting rates of each species and ii) identify complex associations between the environmental conditions and the biting rates.

**Results:** Meteorological and landscape variables were often significantly correlated with the vectors’ biting rates. Many nonlinear associations and thresholds were unveiled by the multivariate models, both for meteorological and landscape variables. From these results, several aspects of the bio-ecology of the main malaria vectors were precised or hypothesized for the Diébougou area, including breeding sites typologies, development and survival rates in relation to weather, flight ranges from breeding sites, dispersal related to landscape openness.

**Conclusions:** Using high resolution data in an interpretable machine-learning modeling framework proved to be an efficient way to enhance the knowledge of the complex links between the environment and the malaria vectors at a local scale. More broadly, the emerging field of interpretable machine-learning has significant potential to help improving our understanding of the complex processes leading to malaria transmission.

## Background

Malaria is a vector-borne disease transmitted by *Anopheles* mosquitoes still affecting 229 million people and causing more than 400 000 deaths worldwide annually (1). Malaria control efforts, mainly through the massive use of long lasting insecticidal nets (2), led to a sustained decrease of the disease burden between 2000 and 2015 (1). However, malaria cases have stalled in the past five years or even increased in specific areas (1). Vector resistance to insecticides, population growth and environmental changes are involved in such worrying trends (3, 4). For effective and sustainable Vector Control (VC), it is now acknowledged that locally tailored interventions, built on a thorough knowledge of the local determinants of malaria transmission, are needed (3–5). To do so, it is of particular importance to decipher with vector bio-ecology at fine and operational spatiotemporal scales (3,4,6) To develop relevant vector control strategies, important features of malaria vectors ecology such as breeding sites typologies, development and survival rates, flight ranges or dispersal have to be considered. Meteorological conditions (temperature, precipitation) and land cover are major environmental factors frequently used to define the ecological niche of malaria mosquitoes (5), in complex, sometimes hardly hypothetizable, ways. Temperature affects the mosquito life history traits, non-linearly, at each stage of its lifecycle (e.g. larval growth, adult survival, biting rate). For example, the daily mortality rate of several adult *Anopheles* species follows a unimodal relationship with air temperature, with an optimal adult survival rate at around 25°C (7–9). Rainfall generates mosquito breeding sites and is therefore an important factor explaining the species seasonality. However, excessive rainfall can destroy developing larvae by flushing them out of their aquatic habitat (10, 11). Land cover may affect mosquito population dynamics by creating breeding sites in hydromorphic lands or altering the dispersal ability of mosquitoes. A modification of land cover/use may therefore either increase or decrease vector abundance relative to species ecological preferences. As an example, deforestation can increase larval breeding sites of malaria vector growing in sunny puddles, whereas it destroys habitats of some deep-forest *Anopheles* species (5, 12). Moreover, even when found together, *Anopheles* species often exhibit specific ecological preferences (13, 14). As an example, *Anopheles gambiae s.s.* was more frequently observed in temporary, rainfall-dependent breeding sites (15–17), whereas *Anopheles coluzzii* showed preferences for more permanent breeding sites (17–19). Altogether, these examples illustrate that vector ecology is finely tuned with environment. Using large scale environnemental indicators could therefore jeopardize the characterization of ecological niches of malaria vectors and consequently lead to sub-optimal or even inappropriate VC intervention at a smaller scale (5). To overcome this issue, we propose to use high resolution Earth Observation (EO) data and develop novel statistical modeling approaches.

Indeed, in malariometric statistical modeling studies, “data” models (20) like linear or logistic regressions are traditionally used with environmental variables extracted from EO data (21–23). These models are well suited for testing pre-established hypotheses about theoretical constructs (e.g. to answer questions like ‘how higher is mosquito abundance for each additional mm of rainfall ?’); however, to explore hypotheses and extract knowledge in complex systems, machine-learning (ML) “algorithmic” models might be more suitable (20, 24). In fact, these models are inherently able to capture complex patterns (such as non-linear relationships and complex interactions between variables) contained in data. After a good predictive algorithm is fitted to a dataset, post-hoc interpretation methods may uncover the complex relationships contained in the data and learned by the model, which can be in turn carefully linked to prior knowledge to identify meaningful - possibly unforeseen - cause-effect relationships, valuable thresholds or interactions (25–27). This predict-then-explain modeling workflow is being increasingly used to generate knowledge from complex datasets (24, 26) and is commonly referred to as “Interpretable Machine Learning” (IML) (25, 26).

In this study, we used a two-step data-driven approach to improve our overall understanding of the ecological niche of malaria vectors in a rural area of southwestern Burkina Faso, where three major malaria vector species are sympatric (29). To do so, we used data from entomological collections carried-out in 27 villages during 15 months and environmental variables (both landscape and meteorological) extracted from EO data. First, we conducted a spatiotemporal correlation analysis to spot out the main ecological factors influencing the human-biting activity of each vector species and get insights on several aspects of their local ecology. In a second stage, we generated algorithmic models (namely Random Forests (28)) that we further analyzed using IML tools to detail the primary and secondary environmental drivers of the human-biting rates, and identify potential complex nonlinear effects and thresholds leading to increased biting risk. We then discussed the environmental drivers of the human-biting activity of the main malaria vectors in the Diébougou area, and briefly concluded with limitations of our work and directions for future research.

## I. Methods

### I.1. Data collection and preparation

#### I.1.1. Entomological data

*Anopheles* human-biting activity was monitored as part of a study carried out in the Diébougou rural health district located in south west Burkina Faso (29). Twenty-seven villages in this 2500 km² wide area were selected according to the following criteria: accessibility during the rainy season, 200–500 inhabitants per village, and distance between two villages higher than two kilometers. Seven rounds of mosquito collections were held in each village between January 2017 and March 2018. The periods of the surveys span some of the typical climatic conditions of this tropical area (3 surveys in the “dry cold” season, 2 in the “dry hot” season, 1 at each extremum of the rainy season) (see Additional figure 1 for the temporal trends of the meteorological conditions).

**Figure 1:**
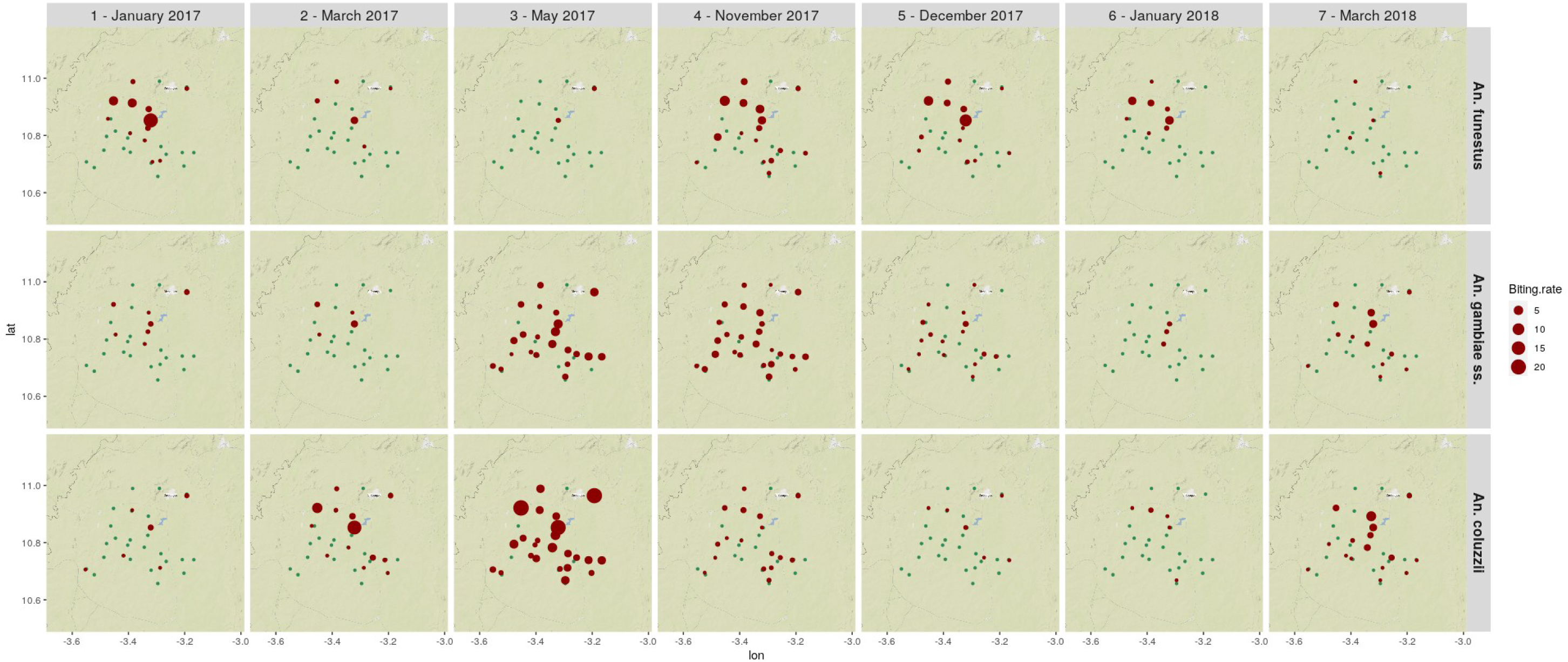
Map of the biting rates of the three main vector species for each village and entomological survey. Unit: average number of bites / human / night. Blue dots indicate absence of bite in the village for the considered survey. Background layer: OpenStreetMap

Mosquitoes were collected using the Human Landing Catch (HLC) technique from 17:00 to 09:00 both indoors and outdoors at 4 sites per village for one night during each survey. The procedure for conducting HLC was for a person to sit on a stool, and mosquitoes to alight on his exposed legs where they were then collected using a hemolysis tube. Collectors were rotated hourly between collection sites and/or position (indoor/outdoor). Independent staff supervised rotations and regularly checked the quality of mosquito collections. Malaria vectors were identified using morphological keys (30, 31). Individuals belonging to the *Anopheles gambiae* complex and the *Anopheles funestus* group were identified to species by PCR (32–34). Mosquito collection design for this study has been described extensively elsewhere (29).

HLC enabled us to measure the presence and abundance of agressive malaria vectors in time and space. In fact, landing on human legs is the behavioral event preceding the biting event. To avoid exposing mosquito collectors to infectious bites, we hence used landing as a proxy for biting, and in turn biting probability / rate as a proxy for the overall presence / abundance of agressive vectors at the time and place of collection.

#### I.1.2. Landscape data

A land cover map of the study area was produced by carrying out a Geographic Object-Based Image Analysis (GEOBIA) (35) using multisource very high and high resolution satellite-derived products. The GEOBIA involved the following main steps: acquisition / collation of the satellite products (Satellite Pour l’Observation de la Terre (SPOT)-6 image acquired on 2017-10-11, Sentinel-2 image acquired on 2018-11-16, and a Digital Elevation Model (DEM) from the Shuttle radar Topography Mission (36)), acquisition of a ground-truth dataset composed of 420 known land-cover samples by both field work (held in November 2018) and photo-interpretation of satellite images, and classification of the land cover over the whole study area using a Random Forest (RF) algorithm (28). The definitions of land cover classes were those proposed by the Permanent Interstate Commitee for Drought Control in the Sahel (37). The resulting dataset was a georeferenced raster image where each 1.5 x 1.5 meter pixel was attributed a land cover class. The confusion matrix was generated using the internal RF validation procedure based on the out-of-bag observations, and the quality of the final classification was assessed by calculating the overall accuracy from the confusion matrix (38).

Spatial buffers were then defined to characterize the environmental conditions at the neighborhood of each HLC collection point. Four buffer radii were considered: 250 meters, 500 meters, 1 kilometer, 2 kilometers. The distance of two kilometers was chosen as the largest radius to minimize overlaps among buffers coming from different villages and because local dispersal of *Anopheles* beyond this distance can be considered negligible (30,39–41) We calculated the percentage of landscape occupied by each land cover class, in each buffer zone, around each collection site. Additional indices related to the presence of water were calculated. The theoretical stream network was produced for the study area, by first generating a flow accumulation raster dataset from the DEM and then applying a threshold value to select cells with accumulated flow greater than 1000 (42). The quality of the product was assessed visually by overlaying it on the SPOT-6 satellite image. We then derived two indices for each collection site: the length of streams in each buffer zone, and the shortest distance to the streams.

In order to describe attractiveness and penetrability of households for malaria vectors, the geographical location of the households in the villages were recorded and two indices were computed. First, the Clark and Evans aggregation index (43) to describe the degree of clustering of the households in each village, as it has been suggested that scattered habitations in a village might increase the attractiveness for some vector species (44). Second, we calculated the distance from each collection point to the edge of the village (defined as the convex hull polygon of each village - i.e. the minimum polygon that encompasses all the locations of the households), as it has been suggested elsewhere that living on the edge can increase biting rates (45).

#### I.1.3. Meteorological data

Daily rainfall estimates were extracted from the Global Precipitation Measurement (GPM) IMERG Final products (46). The raw satellite products were resampled from their original 10 km spatial resolution to a 1 km resolution using a bilinear interpolation method.

Daily diurnal and nocturnal temperatures were derived from the Moderate Resolution Imaging Spectroradiometer (MODIS) Land Surface Temperature (LST) Terra and Aqua products (47, 48). Terra and Aqua daily products were first combined, keeping the highest (respectively lowest) available pixel values for the diurnal (respectively nocturnal) temperature. Missing values in pixels (mostly due to cloud presence) were then filled by temporally interpolating the values of the closest preceding and following available dates.

These meteorological data were collected up to 42 days (i.e. 6 weeks) preceding each mosquito collection, so as to encompass largely the whole duration of the *Anopheles* life cycle in the field (including aquatic and aerial stages) (49). They were then aggregated pixel-by-pixel on a weekly scale (cumulated 7-days rainfall and average 7-days diurnal and nocturnal LST). This temporal granularity represents a reasonable trade-off between the raw, daily information - which might overfit in the statistical models - and larger scales - which might prevent from capturing fine-scale temporal relationships. Next, we calculated the cumulated rainfall and average temperatures for all possible intervals of time available in the data (e.g. b\w 0 and 1 week before the dates of collection, b\w 0 and 2 weeks, b\w 1 and 2 weeks, etc.). The data were finally averaged in the 2 km buffer zone only (considering the 1 km spatial resolution of the source data).

### I.2. Statistical analyses

#### I.2.1. Overall approach

We used a two-step statistical modeling approach to study the relationships between the biting rates of each vector species and the environmental conditions. We first calculated correlation coefficients between the biting rates and the environmental variables at the various buffer sizes / time lags considered. The objectives of this bivariate analysis were twofold: i) to precise some aspects of the ecology of the vectors, and ii) to screen-out variables for the multivariate analysis. In a second stage, we integrated selected variables in multivariate algorithmic models that we further analyzed using interpretable machine learning tools, to seek for potential complex links (nonlinear relationships, relevant thresholds) between the environmental factors and the biting rates.

We ran the whole modeling study separately for each species, as they might exhibit different ecological preferences.

From a statistical point of view, most algorithmic machine learning models, although non-parametric, hardly cope with zero-inflated negative binomial response variables (50, 51), which are typically found in insect count data such as mosquitoes biting rates (52). An alternative approach to model such data is the hurdle model that considers the data responding to two processes: one causing zero *versus* non-zero and the second process explaining the non-zero counts (53). The hurdle methodology in the frame of a widely used algorithmic model (Random Forest), was proposed elsewhere to deal with such distributions of data (51). Besides, this separation is biologically pertinent since it has been shown that the drivers of the presence might differ from those of the abundance (17,44,54). Lastly, it might enable to identify distinct targets for vector control answering to, respectively, eradication (absence of bites) and control (reduction of the number of bites) (17).

We therefore modeled separately the probability of human-vector contact (called ‘presence’ models in the rest of this article) and the positive counts of human-vector contacts (called ‘abundance’ models). Given that HLC data are used as a proxy for human-biting rate, ‘presence’ models analyzed the probability of at least one individual biting a human during a night, while ‘abundance’ models analyzed the number of bites received by one human in one night conditional on their presence (i.e. zero-truncated data). Hence in our ‘presence’ models, the dependent variable was the presence/absence of vectors (binarized as 1/0) collected during 1512 human-nights of HLC (27 villages x 4 collection sites x 2 places (indoors and outdoors) x 7 surveys), while in the ‘abundance’ models, the dependent variable was the number of bites per human during the positive catches sessions - i.e. the sessions with at least one bite.

#### I.2.2. Bivariate analysis using correlation coefficients

The bivariate relationship between the presence / abundance of each vector species and the environmental variables was assessed using multilevel Spearman correlation coefficients (55) with the village entered as a random effect. Multilevel correlations, contrary to simple correlation, account for non-independency between observations in a dataset, by introducing a factor as a random effect in the correlation (on the same principle as random effects in mixed linear regressions).

For each landscape variable, the correlation coefficient was calculated for each buffer radius considered.

For the meteorological variables, Cross-Correlation Maps (CCM) were computed (56) to assess the relationships between the biting rates and the precipitations and temperatures preceding the dates of collection. A CCM enables to study the influence of environmental conditions during time intervals (instead of single time points) prior to the collection event. CCMs hence allow accounting for the effects of accumulated precipitation and average temperature over intervals of weeks preceding the bites (e.g average diurnal temperature between 1 and 3 weeks preceding the bite), instead of single weeks (e.g. average diurnal temperature during the third week preceding the bite).

#### I.2.3. Multivariate analysis using Random Forests and Interpretable Machine Learning

We used the results of the bivariate analysis to select the environmental variables to include as predictors in the multivariate analysis. We first excluded variables that were poorly correlated with the response variable (i.e. correlation coefficients less than 0.1 or p-values greater than 0.2 at all time associations or buffer radii considered), except for variables related to the presence of water - i.e. possible breeding sites - that were all retained whatever their correlation. Then, for each meteorological (resp. landscape) variable, we retained the time lag interval (resp. buffer radius) showing the higher absolute correlation coefficient value. Because the entomological data used in this study were part of a trial, different vector control strategies were implemented in the villages of the study after the third survey. The implemented VC strategies and the place of collection (interior / exterior) were therefore introduced as adjustment variables in our models but their effect on the biting rate was found to be negligible in our analysis and these results will not be discussed further.

Random Forest classifiers were then trained for each species and response variable (presence and abundance models). Random forests are an ensemble machine-learning method that generates a multitude of random decision trees that are then aggregated to compute a classification or a regression (28). They are known for their good predictive capacity, which is mainly due to their ability to inherently capture complex associations between the variables (20). Binary classification RFs were generated for the presence models and regression RFs were generated for the abundance models. The modeling process involved the following steps:

– Feature collinearity: Collinear covariates (i.e. Pearson correlation coefficient > 0.7) were checked for and removed based on empirical knowledge;
– Feature engineering: In the classification models, data were up-sampled within the model resampling procedure to account for the imbalanced structure of the response variable (57). In the regression models, the response variables were log-transformed prior to the model resampling procedure in order to reduce their overdispersion;
– Model training, tuning, selection: Models hyperparameters were optimized using a random 10-combinations grid search (58). For each set of hyperparameters tested, a leave-village-out cross-validation (LVO-CV) resampling method was used. The resampling method involved training the model using in turn the data from 26 of the 27 sampled villages, validating with the data from the remaining village using a predictive performance metric (for the presence model: the Precision-Recall Area Under Curve (PR-AUC); for the abundance model: the Mean Absolute Error (MAE)), and averaging the metric across all hold-out predictions at the end of the procedure. The model retained was the one leading to the highest overall PR-AUC (respectively lowest MAE). The retained model was then fit to all the observations and further used for the interpretation phase;
– Model evaluation: The predictive power of each model was assessed by LVO-CV. We hence evaluated the ability of the models to predict presence or abundance of vectors on unseen human-nights of HLC, whilst excluding from the training sets all the observations belonging to the village of the evaluated observation. Doing so enables to limit overfitting and over-optimistic performance metrics due to spatial autocorrelation (108). For the presence models, Precision-Recall plots were then generated from the observed and predicted values, and the PR-AUC was calculated and compared to the baseline of PR curve (109). For the abundance models, a visual evaluation (i.e. graphical comparison between observed and predicted values) was preferred to a numerical one because performance metrics were expected to be low given the overdispersion of the response data and the type of model used (51). Evaluation plots for the abundance models included i) the distribution of MAEs and ii) observed vs. predicted values for each out-of-sample village.

To interpret the models, we further generated permutation-based Variable Importance Plots (VIPs) (28) and partial dependence plots (PDPs) (59) including standard deviation bands (that can be interpreted as confidence intervals). These plots, part of the interpretable machine learning toolbox, enable to study the effects of one predictor on the response variable while accounting for the effect of the other predictors in the model (25). Variable importance measures a feature’s importance by calculating the degradation of the predictive accuracy of the model after randomly permuting the values of the feature: the higher a variable’s importance, the more that variable contributes to the prediction. Partial dependence plots, on their side, show the marginal effect that one feature has on the predicted outcome (25). PDPs hence help visualize the relationship, learned by a model, between a feature and the response. A PDP is likely to reveal complex (non-linear, non-monotonic) effects when a model has learned such relationships. Importantly, the information provided by these tools should be trusted only if the underlying model has a good predictive power (27).

We finally identified primary and secondary predictors for each model, according to the following criteria. Primary predictors were the top-3 most important predictors of the VIP. Secondary predictors were variables either presenting marked variations in their PDP (e.g. thresholds, significant slopes) or known to influence the bio-ecology of the vector.

##### Software used

The softwares used in this work were exclusively free and open source. The R programming language (60) and the R-studio environment (61) were used as the main programming tools. An R package was developed (111) to extract the NASA meteorological data (MODIS and GPM). The land cover layer was generated using the following R packages: ‘RSAGA’ (62), ‘rgrass7’ (63), ‘raster’ (64), ‘sf’ (65), ‘rgdal’ (66) and ‘randomForest’ (67). The ‘spatstat’ (68) package was used to compute the Clark and Evans aggregation index. The QGIS software (69) was used to create the map of the study area The ‘landscapemetrics’ package (70) was used to calculate the percentage of landscape occupied by each land cover class in the buffer areas. The ‘correlation’ (55) package was used for the correlation analysis. The ‘caret’ (71) and ‘ranger’ (107) packages were used to fit the random forest models in the statistical analysis. The ‘CAST’ (72) package was used to create the temporal folds for cross-validation. The ‘MLmetrics’ (110) package was used to calculate the model evaluation metrics. The ‘iml’ (73) and ‘pdp’ (74) packages were used to generate the partial dependence plots. The ‘patchwork’ (75) package was used to create various plot compositions. The ‘ggmap’ (76) package was used to generate the map of the vectors biting rates. The ‘precrec’ (77) package was used to generate the Precision-Recall plots for the presence models. The ‘tidyverse’ meta-package (78) was used throughout the entire analysis.

## II. Results

### II. 1 Entomological data

A total of 1512 human-nights of HLC were conducted over the twenty-seven villages during the seven entomological surveys. A sum of 3056 vectors belonging to the *Anopheles* genus was collected: 1322 *An. coluzzii*, 708 *An. funestus*, 616 *An. gambiae s.s.*, and 410 from other species. *An. funestus* was present in 12 % of the human-nights of HLC (182 times), while both *An. coluzzii* and *An. gambiae s.s.* appeared in 20 % of the human-nights of HLC (respectively 297 and 302 times). The distribution of the biting rates in the positive sessions (i.e. sessions with at least one bite) was highly left-skewed (for *An. funestus* : median = 2, SD = 4.7, max. = 36; for *An. gambiae s.s.* : median = 2, SD = 1.5, max. = 10; for *An. coluzzii.* : median = 2, SD = 6.8, max. = 50).

Figure 1 shows the distribution of the biting rates of the three main vector species by village and survey. Overall, the map reveals heterogeneous spatiotemporal patterns of biting rates for the three main species. *An. funestus* was found in few villages only, mainly at the end of the rainy season (November) and in the dry-cold season (December, January) (see Additional figure 1 for the temporal trends of the meteorological conditions). It almost disappeared during the dry-hot season (March) and the beginning of the rainy season (May). *An. gambiae s.s.* and *An. coluzzii* were found in almost all the villages at the beginning and the end of the rainy season (May, November). They were also found year-round in some villages, in particular, those located close to dams or the Bougouriba River. *An. coluzzii* was particularly abundant at the beginning of the rainy season (May), while *An. gambiae s.s.* was found in similar abundances at the beginning and the end of the rainy season (May, November).

### II. 2 Land-cover map

Eleven land-cover classes were discriminated: ligneous savannah (52 % of the total surface), crop (25 %), grassland (7 %), marsh (5 %), riparian forest (4 %), woodland (3 %), rice (1 %), settlements (0.5 %), bare soil (0.5 %), main roads (0.3 %), and permanent water bodies (0.3 %). In the buffer areas considered for the modeling study (250 m, 500 m, 1 km, 2 km radii), similar trends were observed regarding the percentage of area occupied by each land cover class (see Additional figure 2). Ligneous savannah included shrub savanna, tree savanna and wooded savanna. Grassland included herbaceous savanna and sahelian short grass savanna. Permanent water bodies included dams and the Bougouriba River. Marshlands included wetland – floodplain and agriculture in shallows and recession. The overall accuracy of the classification was 0.84. The resulting land-cover map of the study area, including the geographical position of the study villages, is presented in Figure 2. Pictures representative of the main land cover classes are provided in Additional file 3.

**Figure 2:**
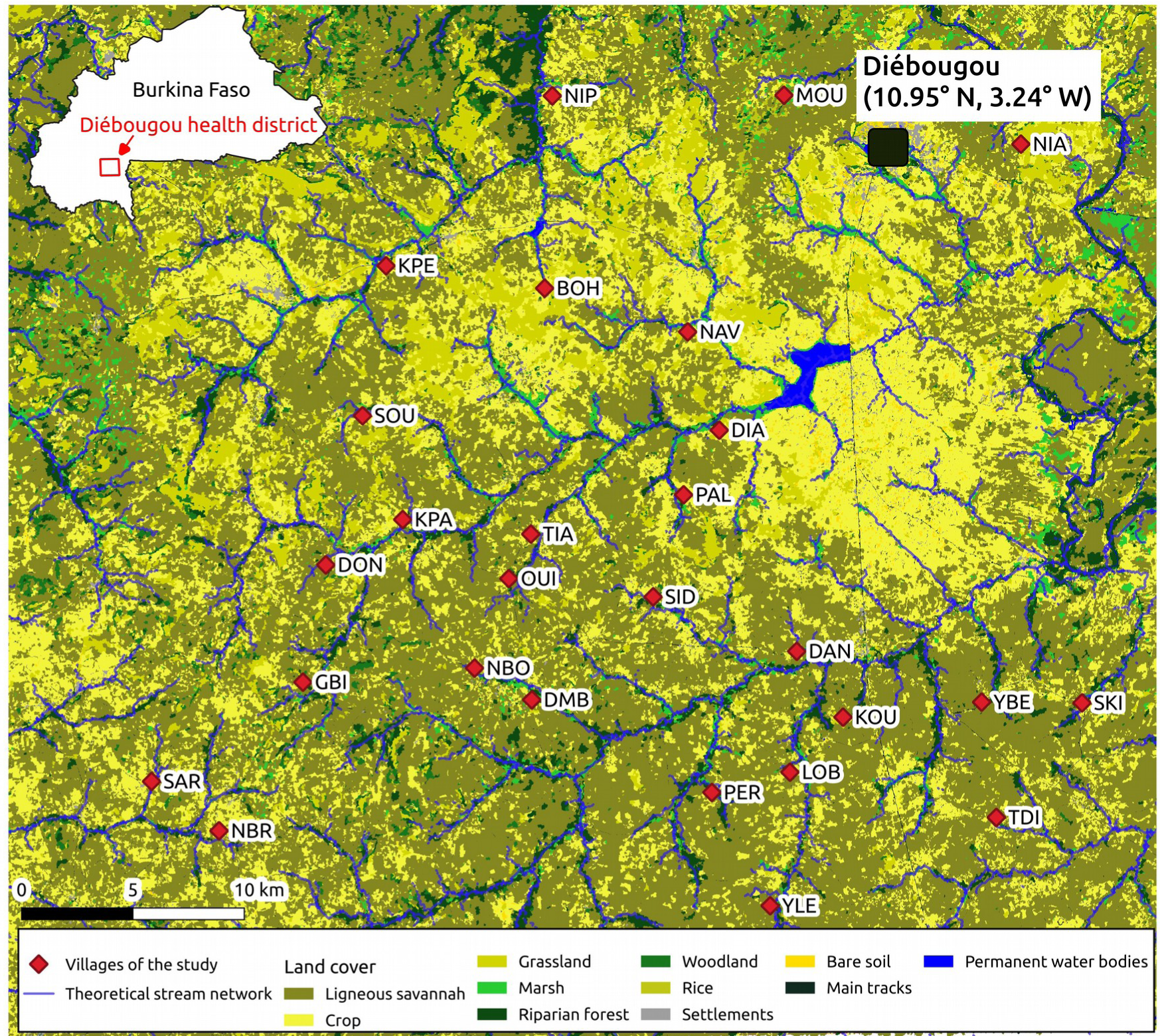
Map of the study area. The map includes the villages of the study, the land cover derived from geographic object-based image analysis of a SPOT-6 satellite image acquired on 2017-10-11 and the theoretical stream network derived from the SRTM DEM.

### II. 3 Bivariate analysis

Figure 3 shows the landscape variables that were significantly correlated (multilevel Spearman’s correlation coefficient (cc) > 0.1 and p-value < 0.2) with the presence or abundance of each of the studied vector species. The presence or abundance of *An. funestus* was correlated to 2 to 6 landscape variables depending to the buffer radius considered, 1 to 5 variables for *An. Gambiae s.s.* and 2 to 5 for *An. coluzzii*. Overall, among the three species, the highest correlation coefficients with the landscape variables were observed for *An. funestus*.

**Figure 3:**
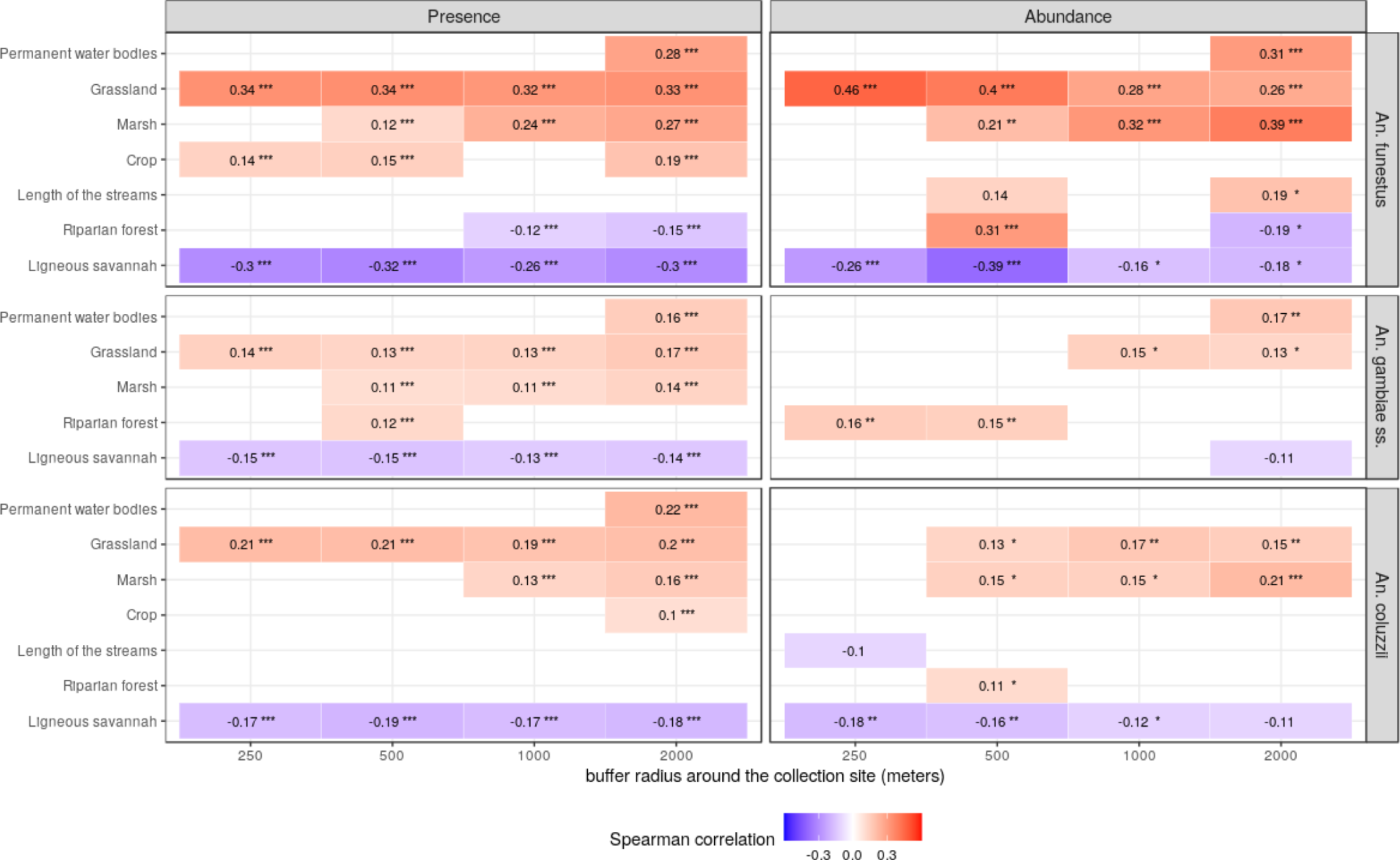
Multilevel Spearman’s correlation between the vectors’ biting rates and the landscape variables. Biting rates were separated into presence / absence of bites (left) and abundance of bites (i.e. positive counts only) (right). Unit of biting rates : nb. landings on human / person / night. Unit of landscape variables: % of landscape occupied by each land cover class. Landscape variables were extracted in 4 spatial buffers zones around the sampling locations (250 m radius, 500 m, 1 km, 2 km) for each main vector species. Only correlations with coefficient > 0.1 and p-values < 0.2 are displayed. Stars indicate the range of the p-value : *** : p-val ∈] 0, 0.001]; ** : p-val ∈ ]0.001, 0.01]; * : p-val ∈ ]0.01, 0.05] : absence of stars : p-val ∈ ]0, 0.001]; ** : p-val ∈ ]0, 0.001]; ** : p-val ∈ ]0.001, 0.01]; * : p-val ∈ ]0.01, 0.05] : absence of stars : p-val ∈ ]0.001, 0.01]; * : p-val ∈ ]0, 0.001]; ** : p-val ∈ ]0.001, 0.01]; * : p-val ∈ ]0.01, 0.05] : absence of stars : p-val ∈ ]0.01, 0.05] : absence of stars : p-val ∈ ]0, 0.001]; ** : p-val ∈ ]0.001, 0.01]; * : p-val ∈ ]0.01, 0.05] : absence of stars : p-val ∈ ]0.05, 0.2[.

Both the presence and the abundance of *An. funestus* were positively correlated with the % of surface occupied by permanent water bodies in the 2-km radius buffer zone. They were also positively correlated with the % of surface occupied by marshlands in all buffer zones with radius >= 500 m, with increasing correlation coefficients as the buffer zone radii increased. The presence and the abundance of *An. funestus* were as well positively correlated with the % of surface occupied by grasslands, and negatively correlated with the % of surface occupied by ligneous savannahs, for all buffer radii. The correlation between the abundance and the % of surface occupied by grasslands and ligneous savannahs increased (both in absolute value and significance) with smaller buffer radii. The presence of *An. funestus* was positively correlated with the % of surface occupied by crops in all buffer zones except for the 1-km radius. The abundance of *An. funestus* was positively correlated with the length of streams in the 250 m and the 2-km radii buffer zones.

Both the presence and the abundance of *An. gambiae s.s.* were positively correlated with the % of surface occupied by permanent water bodies in the 2 km radius buffer zone. Its presence was also positively correlated with the % of surface occupied by marshlands in all buffer zones with radius >= 500 m. The presence and abundance of *An. gambiae s.s.* were as well positively correlated with the % of surface occupied by grasslands (in all buffer zones for the presence, and in the buffer zones with radius >= 1 km for the abundance), and negatively correlated with the % of surface occupied by ligneous savannahs (in all buffer zones for the presence, and in the 2 km radius buffer zone for the abundance). The presence and abundance of *An. Gambiae s.s.* were also positively correlated with the % of surface occupied by riparian forests, only in the 500 m radius buffer zone for the presence, and in buffer zones with radius <= 500 m for the abundance.

The presence of *An. coluzzii* was positively correlated with the % of surface occupied by permanent water bodies in the 2 km radius buffer zone. The presence and the abundance of that species were positively correlated with the % of surface occupied by marshlands in all buffer zones with radius >= 1 km for the presence, and in all buffer zones with radius >= 500 m for the abundance. Presence and abundance of *An. coluzzii* were as well positively correlated with the % of surface occupied by grasslands, and negatively correlated with the % of surface occupied by ligneous savannahs, in all the buffer zones (except in the 250 m radius buffer zone for the abundance). The correlation between the abundance of *An. coluzzii* and the % of surface occupied by ligneous savannahs increased (both in absolute value and significance) with smaller buffer zones.

Figure 4 shows the meteorological variables that were significantly correlated (Multilevel Spearman correlation coefficient (cc) > 0.1 and p-value < 0.2) with the presence or abundance of each of the studied vector species (in the form of cross-correlation maps). Overall, among the three species, the highest correlation coefficients with the meteorological variables were observed for *An. coluzzii,* closely followed by *An. gambiae s.s*.

**Figure 4:**
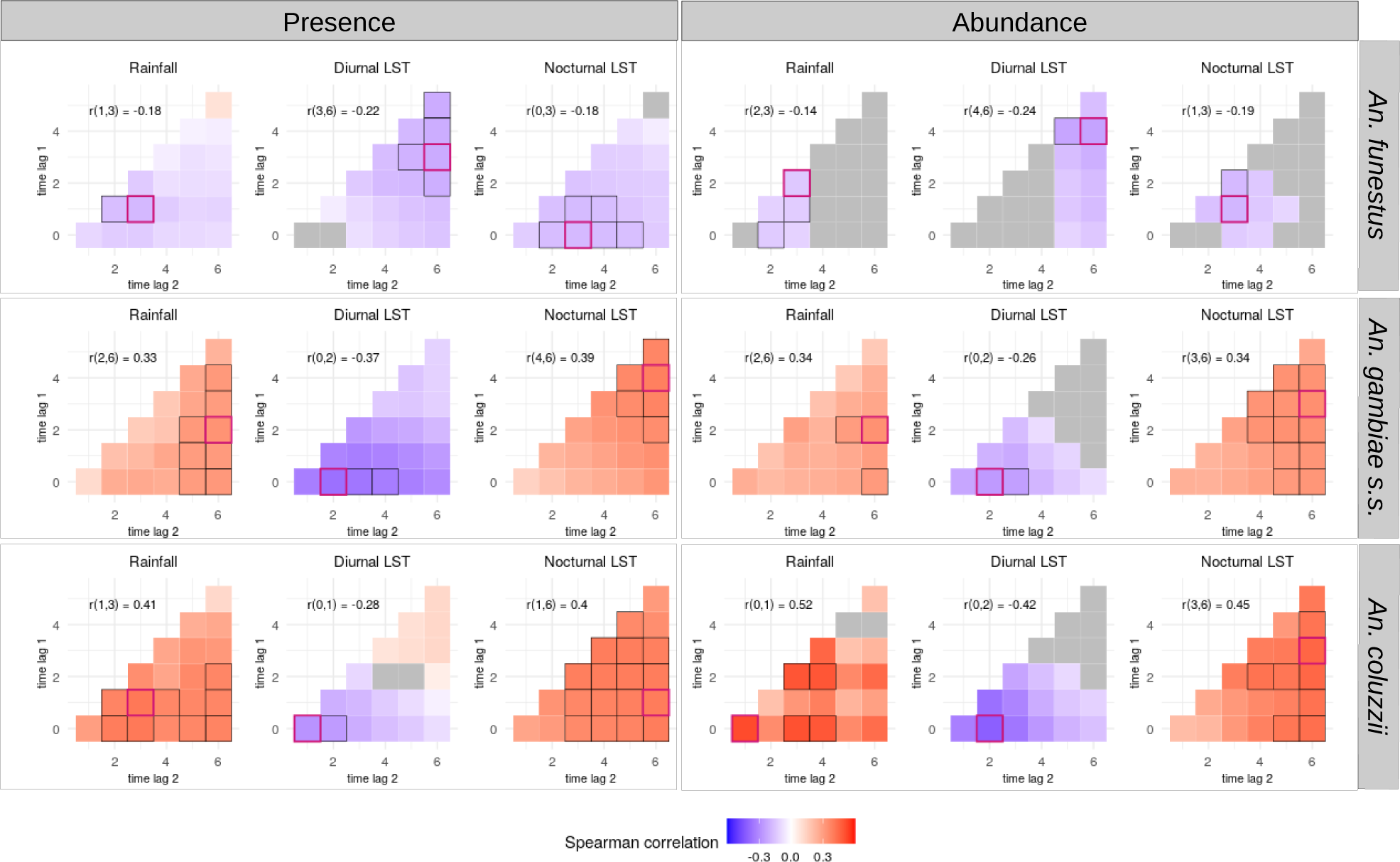
Multilevel Spearman’s correlation between the vectors’ biting rates and the meteorological variables (as cross-correlation maps). Biting rates were separated into presence / absence of bites (left) and abundance of bites (i.e. positive counts only) (right). Unit of biting rates : nb. landings on human / person / night. Unit of meteorological variables: °C for Land Surface Temperatures (LST), cumulated millimeters for rainfall. Meteorological variables were extracted on a weekly scale up to 6 weeks before the dates of collection for each main vector species. In each CCM, time lags are expressed in week(s) before the date of collection. The red-bordered square indicates the time lag interval that showed the highest correlation coefficient (absolute value) with the meteorological variable (the associated time lags interval and correlation coefficient are reported on the top-left corner of the CCM). The black-bordered squares indicate correlations close to the highest observed correlation (i.e. less than 10% of difference). Grey-filled squares indicate correlations with p-value > 0.2 or coefficient > 0.1.

The presence and abundance of *An. funestus* showed quite weak correlations with all the meteorological variables (Spearman’s correlation coefficient always < 0.25) when compared to *An. gambiae s.s. or An. coluzzii*. Correlations between both response variables (presence and abundance) and the three meteorological variables (cumulated rainfall, diurnal LST, nocturnal LST), when significant, were negative. The maximum correlation coefficients between each meteorological variable and both presence and abundance were found: for cumulated rainfall recorded b\w 1-2 and 3 weeks before the date of collection, for diurnal temperatures recorded b\ w 3-4 and 6 weeks before the date of collection, and for nocturnal temperatures recorded b\w 0- 1 and 3 weeks before the date of collection.

The presence and abundance of *An. gambiae s.s.* were positively correlated with cumulated rainfall, at all time lags. The maximum correlation coefficients with both presence and abundance were found for cumulated rainfall recorded b\w 2 and 6 weeks before the date of collection. The presence and abundance of *An. gambiae s.s.* were also positively correlated with nocturnal temperatures at all time lags, the maximum correlation coefficient with both response variables were found for temperatures recorded b\w 3-4 and 6 weeks before the date of collection. The presence and the abundance of *An. gambiae s.s.* were negatively correlated with diurnal temperatures preceding the date of collection at almost all time lags. The maximum correlation coefficient with both response variables was found for temperatures recorded b\w 0 and 2 weeks before the date of collection.

The correlations between meteorological variables and both the presence and abundance of *An. coluzzii* exhibited similar trends as *An. gambiae s.s.*, with few, notable, differences. The presence and abundance of *An. coluzzii* were positively correlated with cumulated rainfall preceding the date of collection at all time lags. The maximum correlation coefficient with cumulated rainfall was found b\w 1 and 3 weeks before the date of collection for presence, and b\w 0 and 1 week before the date of collection for abundance (the correlation coefficient b\w 1 and 3 weeks was also among the highest for the abundance). The presence and abundance of *An. coluzzii* were positively correlated with nocturnal temperatures at all time lags with maximum correlation coefficients found b\w 1/3 and 6 weeks before the date of collection for both response variables. The presence and abundance of *An. coluzzii* were, overall, negatively correlated with diurnal temperatures preceding the date of collection. The maximum correlation coefficient between diurnal temperatures and both response variables was found b\w 0 and 1/2 weeks before the date of collection.

### II. 4 Multivariate analysis

The PR-AUC of the presence models were 0.56 (baseline = 0.12), 0.46 (baseline = 0.20) and 0.60 (baseline = 0.20) for *An. funestus*, *An. gambiae s.s.*, and *An. coluzzii* respectively, indicating good predictive accuracies of the models. These models often slightly overestimated low presence probabilites (i.e. high number of false positives). The abundance models reflected the trends well for the three species, although they often underestimated high counts. The models evaluation plots are available in Additional figures 4 and 5 (for the presence models : Precision-Recall plots and observed vs. predicted values for each out-of-sample village; for the abundance models : distribution of MAE and observed vs. predicted values for each out-of-sample village). Figures 5, 6 and 7 show the model interpretation plots (variable importance plot and partial dependence plots) for *An. funestus*, *An. gambiae s.s.*, and *An. coluzzii,* respectively.

**Figure 5:**
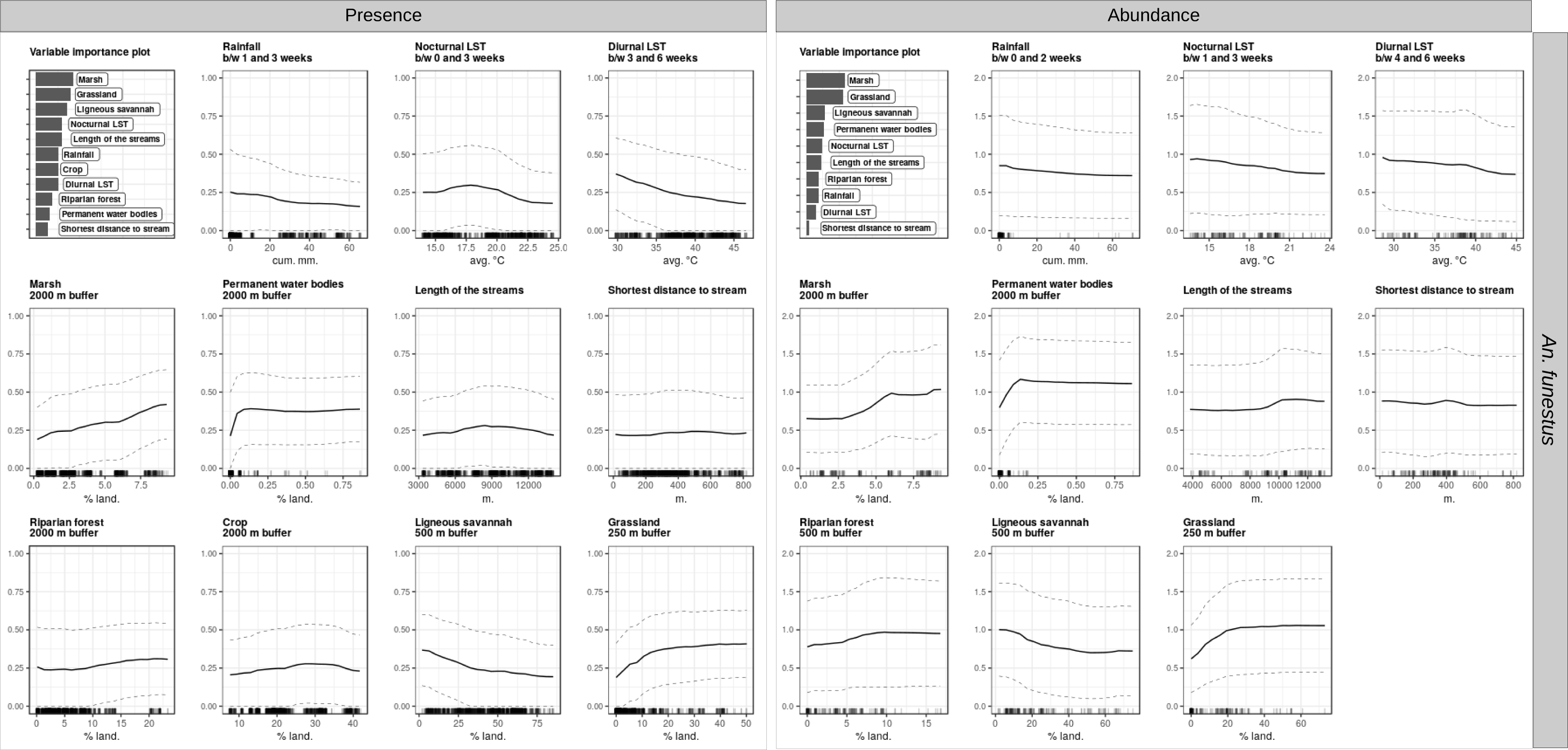
Interpretation plots of the Random Forest models for An. funestus. Biting rates were separated into presence / absence of bites and abundance of bites (i.e. positive counts only) and two models were therefore generated (presence - left - and abundance - right). For each model, the top-left corner plot is the variable importance plot. The other plots are partial dependence plots (PDPs) for each variable included in the models (1 plot / variable). The y-axis in the PDPs represents: in the presence models, the probability of at least one individual biting a human during a night; in the abundance models, the log-transformed number of bites received by one human in one night conditional on their presence. The dashed lines represent the partial dependence function + / - one standard deviation (i.e. variability estimates). The range of values in the x-axis represents the range of values available in the data for the considered variable. The rugs above the x-axis represent the actual values available in the data for the variable. LST stands from “Land surface temperature”.

**Figure 6:**
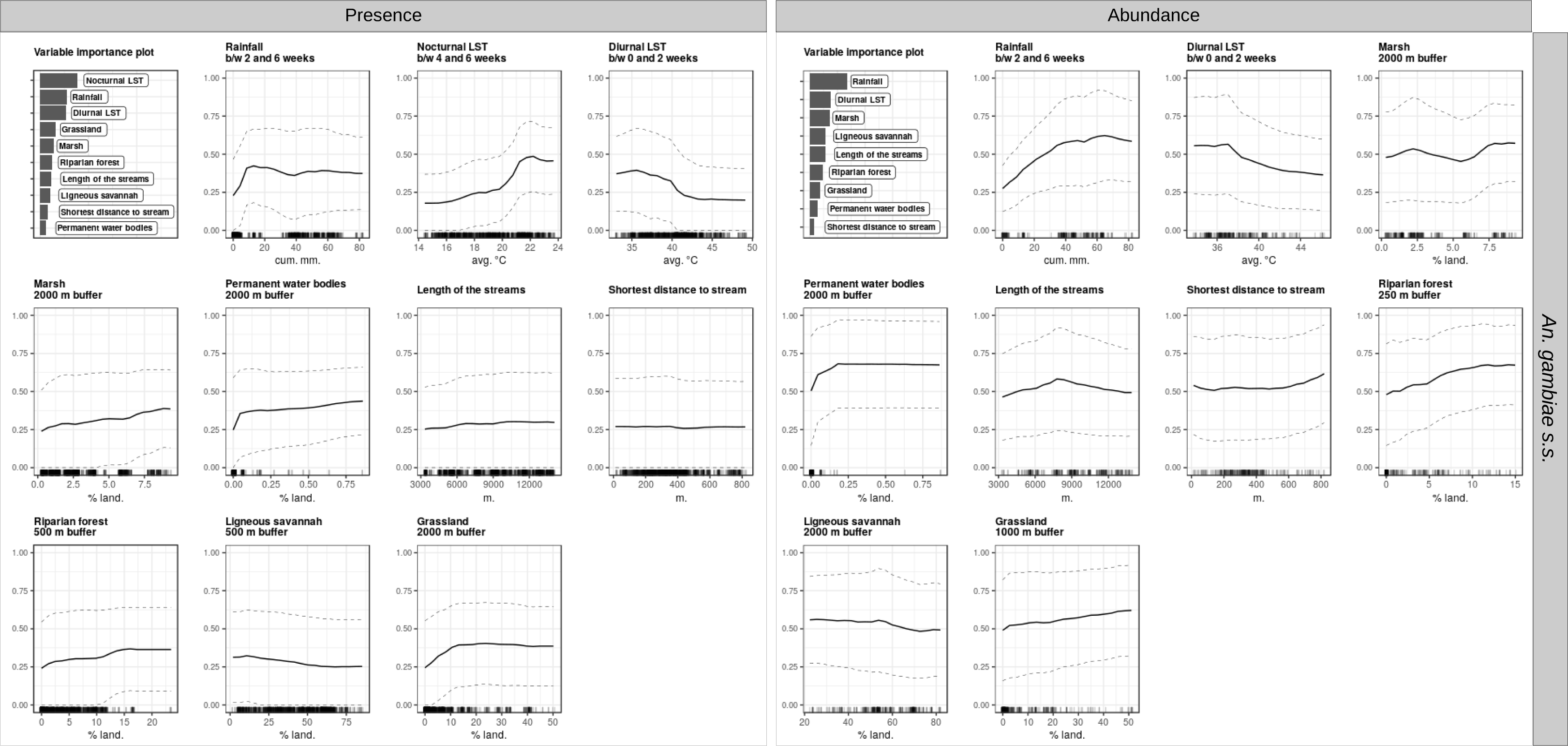
Interpretation plots of the Random Forest models for An. gambiae s.s. Biting rates were separated into presence / absence of bites and abundance of bites (i.e. positive counts only) and two models were therefore generated (presence - left - and abundance - right). For each model, the top-left corner plot is the variable importance plot. The other plots are partial dependence plots (PDPs) for each variable included in the models (1 plot / variable). The y-axis in the PDPs represents: in the presence models, the probability of at least one individual biting a human during a night; in the abundance models, the log-transformed number of bites received by one human in one night conditional on their presence. The dashed lines represent the partial dependence function + / - one standard deviation (i.e. variability estimates). The range of values in the x-axis represents the range of values available in the data for the considered variable. The rugs above the x-axis represent the actual values available in the data for the variable. LST stands from “Land surface temperature”.

**Figure 7:**
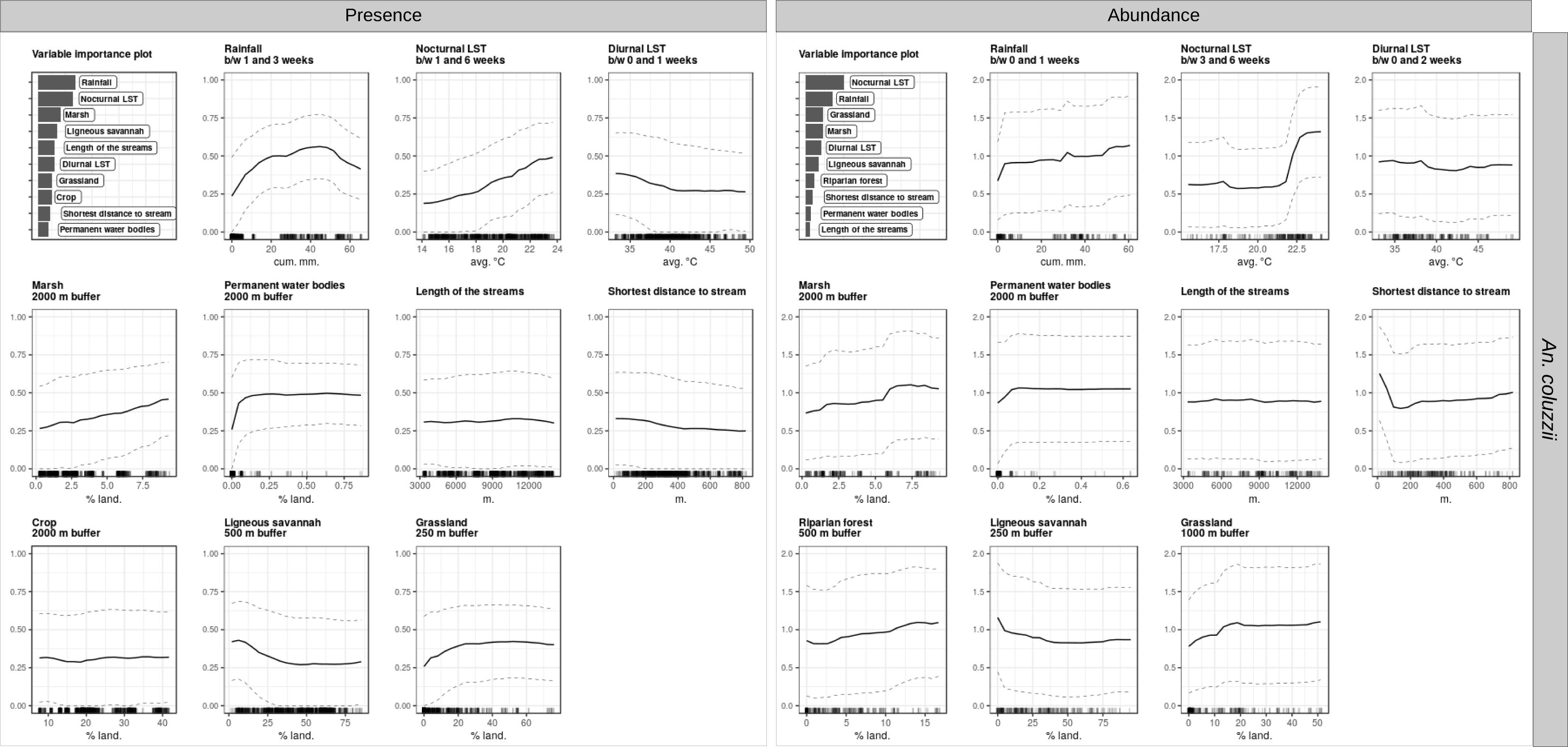
Interpretation plots of the Random Forest models for An. coluzzii. Biting rates were separated into presence / absence of bites and abundance of bites (i.e. positive counts only) and two models were therefore generated (presence - left - and abundance - right). For each model, the top-left corner plot is the variable importance plot. The other plots are partial dependence plots (PDPs) for each variable included in the models (1 plot / variable). The y-axis in the PDPs represents: in the presence models, the probability of at least one individual biting a human during a night; in the abundance models, the log-transformed number of bites received by one human in one night conditional on their presence. The dashed lines represent the partial dependence function + / - one standard deviation (i.e. variability estimates). The range of values in the x-axis represents the range of values available in the data for the considered variable. The rugs above the x-axis represent the actual values available in the data for the variable. LST stands from “Land surface temperature”.

The most important predictors of the presence and abundance of *An. funestus* were landscape-based, including: % of surface occupied by marshlands (in the 2 km radius buffer zone), grasslands (in the buffer zones radii <= 500 m), and ligneous savannahs (in the 500 m radius buffer zone). The probability of presence of *An. funestus* increased linearly with surface occupied marshlands in the range available in the data (0% - 10%), while the abundance was constant in the range 0% - 3%, increased approximately linearly from 3% to 6%, and finally stabilized in the range 6% - 10%. Both the probability of presence and the abundance increased linearly with surface occupied by grasslands in the range 0% - 20%, and stabilized above that threshold. Conversely, they decreased linearly with surface occupied by ligneous savannahs in the range 0% - 50%, and above that threshold stabilized for abundance and tend to diminish (with a lower trend though) for presence.

Secondary predictors of the presence and abundance of *An. funestus* were : % of surface occupied by permanent water bodies in the 2 km radius buffer zone (increase in the range 0% - 0.1%, stable in the range 0.1% - 1%), diurnal LST b\w 3 to 4 weeks and 6 weeks before the date of collection (negative, approximately linear, association), and nocturnal LST b\w 0 to 1 and 3 weeks before the date of collection (stable in the range 14°C – 19°C, decrease in the range 19°C - 25°C).

The most important predictors of the presence of *An. gambiae s.s.* were meteorological variables, namely: nocturnal LST (b/w 4 and 6 weeks, i.e. 28 and 42 days, before the date of collection), rainfall (b/w 2 and 6 weeks before the date of collection), and diurnal LST (b/w 0 and 2 weeks before the date of collection). The probability of presence of *An. gambiae s.s.* increased slowly for nocturnal LSTs in the range 14°C - 20°C and more rapidly above that threshold. It increased linearly with the cumulated rainfall in the range 0 mm - 10 mm, and was stable above that threshold. It decreased for diurnal LSTs in the range 35°C - 41°C, and stabilized above that threshold. Secondary predictors of the presence of *An. gambiae s.s.* were: % of surface occupied by grasslands in the 2 km radius buffer zone (increase in the range 0% - 15%, stable above), % of surface occupied by ligneous savannahs in the 500 m radius buffer zone (negative linear association), % of surface occupied by marshlands in the 2 km radius buffer zone (positive linear association).

When *An. gambiae s.s*. was present, accumulated rainfall (b/w 2 and 6 weeks before the date of collection) was, by far, the most important predictor of its abundance. Other primary predictors were diurnal temperatures (b/w 0 and 2 weeks before the date of collection) and % of surface occupied by marshlands (in the 2 km radius buffer zone). The abundance of *An. gambiae s.s.* increased linearly in the range 0 mm - 50 mm of cumulated rainfall and stabilized above that threshold (range 50 mm – 80 mm). It slowly increased with the % of surface occupied by marshlands. Secondary predictors of the abundance of *An. gambiae s.s.* included : % of surface occupied by ligneous savannahs in the 2 km radius buffer zone (negative linear association), % of surface occupied by permanent water bodies in the 2 km radius buffer zone (increase in the range 0% - 0.1%, stable in the range 0.1% - 1%), % of surface occupied by riparian forests in the 250 m radius buffer zone (positive linear association), and % of surface occupied by grasslands in the 1 km radius buffer zone (positive linear association).

The most important predictors of the presence of *An. coluzzii* were: accumulated rainfall (b/w 1 and 3 weeks before the date of collection), nocturnal LST (b/w 1 and 6 weeks before the date of collection) and % of surface occupied by marshlands (in the 2 km radius buffer zone). A total of approximately 40 mm of rainfall b/w 1 and 3 weeks before the date of collection was enough to double the probability of presence of *An. coluzzii* (from an average 0.25 without rainfall to 0.55). Beyond that amount of rainfall, the probability of presence tended to diminish (range 50 mm – 65 mm). The probability of presence of *An. coluzzii* increased linearly with nocturnal LSTs and % of surface occupied by marshlands. Secondary predictors of the presence of *An. coluzzii* included : diurnal LST b/w 0 and 1 week before the date of collection (decrease in the range 34°C – 40%, stable in the range 40°C-50°C), % of surface occupied by ligneous savannahs in the 500 m radius buffer zone (decrease in the range 0% - 40%, stable above), % of surface occupied by grasslands in the 250 m radius buffer zone (increase in the range 0% - 30%, stable above), % of surface occupied by permanent water bodies in the 2 km radius buffer zone (increase in the range 0% - 0.1%, stable in the range 0.1% - 1%).

When *An. coluzzii* was present, primary predictors of its abundance were nocturnal LST (b/w 3 and 6 weeks before the date of collection), cumulated rainfall (b/w 0 and 1 week before the date of collection), and % of surface occupied by grasslands (in the 1 km radius buffer zone). The abundance of *An. coluzzii* was constant for nocturnal LSTs under 22°C and strongly increased above, until 23°C. The association between abundance and cumulated rainfall was quite weak, but overall positive. The abundance increased linearly with the surface of grasslands in the range 0% - 20%, and stabilized above that threshold. Secondary predictors of the abundance of *An. coluzzii* were : % of surface occupied by marshlands in the 2 km radius buffer zone (positive linear association), % of surface occupied by riparian forests in the 500 m radius buffer zone (positive linear association), % of surface occupied by ligneous savannahs in the 250 m radius buffer zone (decrease in the range 0% - 40%, stable above), and distance to the closest stream (decrease in the range 0 m - 100 m, stable above).

Notably, the confidence intervals of the partial dependence functions were overall high for all species and variables and, with a few exceptions, no variable emerged as much more predictive than others (in the VIPs) nor had signals outstandingly strong (in the PDPs).

## III. Discussion

In this modeling study, we linked the biting rates of three major malaria vector species with environmental conditions at vicinities of places and periods of times of biting events to better understand their bio-ecology at fine spatio-temporal scales and identify important factors leading to increased biting risk. First, we correlated the biting rates of the vector species with i) each meteorological variable at various time lags before the mosquito collection (using cross-correlation maps) and ii) each landscape variable in various buffer zones around the HLC locations. Then, for selected time lags or spatial radii (the ones with the highest correlation coefficients), we generated multivariate models to study i) the contribution of each environmental variable in predicting the biting rates and ii) the nature of the relationship between each environmental variable and the biting rates (all other environmental conditions considerated).

In this section, we first discuss the relationships between the biting rates of the malaria vectors and the meteorological and landscape conditions in the Diébougou area, and link them to the bio-ecology of the species. We then briefly summarize some of the advantages of the modeling method used for knowledge generation in the field of landscape entomology. We conclude the discussion with some limitations of this study and directions for future research.

### III.1 Effects of meteorological variables

The cross-correlation maps enable to study how meteorological conditions affect the various stages of the mosquito lifecycle (56). Here, we found that weather conditions (rainfall, nocturnal LST and diurnal LST) were significantly correlated with, and almost always primary predictors of, the presence and abundance of the species of the *Anopheles gambiae* complex. Stronger effects of these meteorological variables were found at various time lags in the studied range (from 0 to 6 weeks before collections). As discussed by Lebl and colleagues (79), weather dependent life expectancy and development rates make it difficult to link time lags (of weather recordings) influencing mosquito abundances to different development stages. Given the mean lifespan and larval development duration of the *Anopheles* species collected in our area (49,80,81), weather during the first week (i.e. b\w 0 and 1 week) before collection was expected to influence adult lifetime of collected mosquitoes, and weather during weeks 1-3 (i.e. b\w 1 and 2-3) before collection was expected to influence larval lifetime of collected mosquitoes. Weather during preceding weeks (i.e beyond 3 weeks before the date of collection) might affect observed densities by influencing survival and development rates of i) parent generations through mechanical effect on the population dynamic (79), ii) the current / sampled generation through maternal / paternal effects (82, 83), or iii) the current generation by preparing different biotic and abiotic conditions (for instance, by filling suitable larval development sites with water or by allowing the development of food sources, competitors or predators of *Anopheles* larvae).

In the spatiotemporal frame of our study, nocturnal LST ranged from 14°C to 24°C, and diurnal LST from 33°C to 50°C. Both nocturnal and diurnal LST were important predictors of the presence and abundance of *An. gambiae s.s. and An. coluzzii*, often with marked thresholds. Indeed, for *An. gambiae s.s.* we were able to identify a threshold of minimal LST over which the probability started to increase (20°C), and for both species we also identified a threshold of maximum LST over which the probability reached a minimum (40°C). For both species, diurnal temperature had their strongest effect during the 2 weeks preceding the dates of collection. This indicates that increasing diurnal temperatures probably reduced adult survival and larvae development rates of the sampled generation of mosquitoes. Indeed, high temperatures are known to inhibite development of anophelines larvae (84) and to reduce adult survival (85). Regarding nocturnal temperatures, the time period with the strongest effect on presences and abundances of both *An. gambiae s.s.* and *An. coluzzii* was comprised between 3 and 6 weeks before collection. This indicates that nocturnal (i.e. minimal) temperatures had their strongest impact by either affecting previous generations (low temperature are known to reduce adult survival and inhibite larval development (84, 85)) or modifying habitats (with a delay) for the collected generation (for instance, low temperatures may inhibit the development of algae (86) which biomass was found to be associated with larval densities (87, 88)).

The high correlation coefficients between cumulated rainfall and both the presence and abundance of *An. gambiae s.s.* and *An. coluzzii* and the fact that rainfall was systematically an important predictor of these species, might indicate that in our area *An. gambiae s.s.* and *An. coluzzii* are preferably attached to rainfall-dependent breeding sites, confirming other studies (19,89,90) and explaining their seasonality. The time period with the strongest effects of rainfall on presence and abundance of *An. gambiae s.s.* was comprised between 2 and 6 weeks before collection, suggesting an effect on parental generations (as observed by Lebl and colleagues (79) for other mosquito species) or by modifying habitats (abiotic and/or biotic conditions) for the collected generation of these species. Conversely, rainfall had one of its highest correlation coefficient with the presence and abundance of *An. coluzzii* during weeks 1-3 before the dates of collection, indicating that rainfall might had their greatest influence on the larval stages of the sampled generation of mosquitoes.

Different amounts of rainfall were needed for *An. coluzzii* and *An. gambiae s.s.* to be present or abundant, suggesting different breeding habitats preferences. Few rainfall was needed for *An. gambiae s.s.* to increase its probability of being present; additionally, rainfall was by far the most important predictor of its abundance (with a strong positive and approximately linear association). This could indicate that *An. gambiae s.s.* was more attached to breeding sites that quickly appear (presence) and abound (abundance) when few rain falls and disappear after it stops, i.e. temporary breeding sites like puddles. *An. coluzzii* needed more rainfall to be present, suggesting preferences for breeding sites that require more water to be flooded, i.e. semi-permanent surface water collections like marshlands or streams, which are usually filled-in by rainfall throughout the rainy season and shortly after. Indeed, the % of surface occupied by marshlands was significantly correlated with, and the fourth most important predictor of, the abundance of *An. coluzzi*. Overall, these hypotheses about the preferred breeding habitats of *An. gambiae s.s.* and *An. coluzzii* confirm the literature (15–19).

### III.2 Effects of landscape variables

The biting rates of the three species were significantly correlated with several landscape variables, at varying distances from the collection points, and with fluctuating correlation coefficients. In addition, primary predictors of the presence and abundance of *An. funestus* were systematically landscape-based, and some were also primary predictors of the abundance of *An. gambiae s.s.* and *An. coluzzii*. Overall, this indicates that local landscape conditions are important drivers of the bio-ecology of the malaria vectors in rural areas, confirming the literature (5,12,17).

The mere presence of permanent water bodies (irrespective of the surface that they occupied) was sufficient to increase (even moderately) the probability of presence and the abundance of the three species. Permanent water bodies, where available, are likely to form breeding habitats for the *Anopheles* species (17,44,91–93), and explain why the few study villages located close to the dams and the main river are exposed to year-round bites of high densities of the three species (29). The % of surface occupied by marshlands at the vicinities of the biting sites was the most important predictor of the presence and abundance models of *An. funestus*. In our study area, marshlands, a semi-permanent aquatic environment, hence seemed to be one of the preferred breeding habitat of *An. funestus*, as it has been observed in other places (91, 92). Notably, the correlation coefficients between the presence / abundance of bites and the % of surface occupied by breeding habitats land cover types (marshlands and permanent water bodies) increased as buffer sizes increased. This might indicate that mosquitoes are able to fly over quite large distances to reach their biting site from these breeding habitats (>= 2 km), as observed elsewhere in similar landscapes (94, 95). Proximity to the streams (< 100 m) and % of landscape occupied by the riparian forests (<= 500 m) were secondary predictors of the presence and/or abundance of *An. gambiae s.s.* and *An. coluzzii*. Streams and riparian forests (which are spatially interrelated, i.e. streams flow under riparian forests) might hence form secondary, semi-permanent, breeding sites for the species of the *Anopheles gambiae* complex in the Diébougou area.

Grasslands - a very “open” landscape - and ligneous savannahs - the most “closed” landscape in our study area – were alone or together, significantly correlated with, and important predictors of, the presence and/or the abundance of the three malaria vector studied. Increasing surfaces of grassland areas were associated with increasing probabilities of presence or abundances, while increasing surfaces of savannahs were associated with decreasing probabilities of presence or abundances. With some rare exceptions, these landscape indicators were most highly correlated in small radii (<= 500 m) buffer areas around the collection sites. Although grassland may provide suitable breeding sites for, at least, *An. gambiae s.s.* and *An. coluzzii* (96), these results seems to indicate that the degree of openness of the landscape some hectometers around villages had a great impact on malaria mosquitoes biting rates. Our observations are supported by the hypothesis of the lower dispersal of *Anopheles* mosquitoes in closed landscape (in comparison to open landscape) (97) leading to shorter gonotrophic cycles durations and therefore increased biting frequencies and higher biting rates (98). A similar observation was previously made with *An. coluzzii* in Benin (17). In the Diebougou area, ligneous savannahs seemed to act as natural protective barriers against the malaria vectors and conversely, grasslands as an aggravating biting risk factor. In a country which is increasingly replacing its closed landscapes (savannahs) by opened ones (37), this observation is worrying for malaria transmission. Removal of savannahs may significantly increase biting densities. This concern may however be mitigated for our study area, as savannahs are usually mainly replaced by crops (37), which themselves did not seem to be an aggravating risk factor (i.e. crops did not come out as an important variable in our models).

### III.3 Methodological bonus : on the use of algorithmic models and interpretable machine learning in landscape entomology

In our study, we have shown how complex statistical models and IML can be used to enhance the fundamental knowledge and understanding of the complex links between the environment and the malaria vectors. Advantages of this modeling workflow over more traditional modeling methods (e.g. linear of logistic regressions) include the ability to i) inherently capture and unveil complex patterns such as non-linear or non-monotic relationships (e.g. effect of temperature and rainfall) and interactions and ii) easily include more variables (20) and hence capture small – yet relevant - effects (e.g. effects from riparian forests or distance to streams). Necessary conditions to perform causal interpretation from “black-box” models are to i) generate a good predictive model and ii) have some prior domain knowledge about the causal structure of the system under study (27). Both conditions were met in our work.

Using machine-learning black-box models for scientific discovery, i.e. to generate new knowledge, from data, is an emerging trend in many disciples (26, 99) that has been made possible by the recent development of both IML tools (73) and the know-how to interpret these complex models (25–27,99,100). ML models enable to integrate knowledge from existing theory in a less formal way than “data” models, and as such can be useful for theory development, provided that a careful linkage to existing knowledge is made (99, 101). New theoretical insights generated from data and models may then in turn lead to unforeseen experimental research questions. We believe that the fields of landscape epidemiology and entomology still need to fully embrace the potential offered by these methods, in support not only of prediction or forecasting, but also explanation, i.e. to improve our understanding of the complex processes leading to malaria transmission.

### III.4 Limitations and directions for future research

An important limitation of our work is linked to the spatio-temporal sampling distribution of mosquito collection. First, no collection was conducted during the high rainy season (July to October), at the known mosquito abundance and malaria transmission peaks. Second, all the collection points were less than 800 m away from the theoretical hydrographic network (which is spatially interrelated with many breeding habitats such as marshlands, streams, riparian forests), meaning that our study could not identify potential differences in the drivers of vector abundances for households further than this distance. Year-round longitudinal collections, including sites further away from permanent or semi-permanent breeding habitats, may enable to better understand of the overall malaria mosquito spatio-temporal dynamics in the area. Meanwhile, these limitations must be accounted for if predictive maps of the spatio-temporal distribution of vector abundances are generated based on our models.

Another limitation is the nature and diversity of the variables introduced in the models. Very fine scale potential important drivers of mosquito abundance, such as the presence of alternative sources of blood-meal (e.g. cattle), of domestic breeding sites, market gardening, or the micro- climatic conditions on the night of collection, have not been investigated. These variables were significant drivers elsewhere in West African rural settings (17,44,102). Yet, the good predictive accuracies of the models suggest that, most probably, the most important drivers of vector abundance our study area have been identified.

The absence of a strong signal from single variables in the models, as well as the large confidence intervals in the PDPs, suggest that the models might have learned important interactions between variables. IML tools such as the H-statistic (103) might help reveal such interactions, and others like the 2-variables PDP (73) might help explore their effect on malaria vectors abundances. Other tools can be used to analyze individual, or a target set of, predictions made by the models (these tools are called *local interpretation methods*, e.g. LIME (104) or Shapley values (105)). Local interpretation could be useful e.g. to precise the environmental drivers of vector presence / abundance for a village of interest, or to better spot- out the drivers of the spatial heterogeneity of biting rates within a season of interest. Altogether, these IML tools might enable dig deeper into the models and hence, the complexity of the ecological niche of malaria vectors.

This work has revealed how landscape can influence the biting rates of the vectors. Further investigations on the role played by the level of openness of the landscape are needed to confirm the hypotheses rose. A finer-grained land cover classification (e.g. discriminating shrub, tree and wooded savanna) could help precise some of our hypotheses: for instance, is there a correlation between the gradient of closedness of the savannahs and the abundance of vectors in our study area? Moreover, additional field work could help precise the potential cause-effect relationship between surface of grasslands and malaria vector presence / abundance (i.e. breeding habitat or open landscape favoring dispersal?).

Lastly, the good predictive accuracies of our models lays the ground for the development of operational tools to support vector surveillance and improve forecasting of epidemics outbreaks at local scale. Indeed, in this study we were able to build models predicting the biting densities in space and time, with good predictive powers, although prediction was not the main purpose. Using additional variables extracted from EO data, creating composite variables (e.g. NDVI), or conducting a spatiotemporal feature selection (108), are all techniques that might improve the performance of our spatio-temporal models. Such predictive models could in turn be the cornerstone of an early warning system that forecast the spatiotemporal distribution of vector biting rates at fine spatiotemporal scales.

## Conclusion

In this study, several aspects of the bio-ecology of the main malaria vectors in the Diébougou area were explored using field mosquito collections and high resolution EO data (traducing both meteorological and landscape local conditions) in a state-of-the-art statistical modeling framework. Overall, the spatio-temporal distributions of biting rates of *An. coluzzii* and *An. gambiae s.s.* were tightly associated to meteorological conditions (temperature, precipitation), while those of *An. funestus* were more closely linked to landscape conditions. Meteorological conditions (temperatures, rainfall) putatively affected all developmental stages of the mosquitoes (larval, adult), at varying levels according to the species and the meteorological variable. Weather occasionally had an even greater impact on time periods preceding the lifespan of the sampled generation. Primary and possible secondary breeding habitats of each vector species were proposed: *An. funestus*, *An. coluzzii*, and *An. gambiae s.s.* seemed to be distributed along a gradient of persistence of the breeding sites, from permanent to temporary, confirming the literature (17,90,106). The rate of openness of the landscape seemed to play a major role in the biting rates, which could represent a major concern in a context of changing landscapes. This work lays the foundations for the development of early warning systems for the detection of malaria outbreaks at fine spatiotemporal scales.

## Additional files

Additional figure 1 (add_fig_1.png): Summary of the meteorological conditions around the sampling points. Average meteorological conditions in a 2 km radius buffer zone around the collection points (weekly aggregation). Vertical red lines indicate the dates of the entomological surveys. Ribbons indicate the mean +/- one standard deviation considering all the sampling points for the week.

Additional figure 2 (add_fig_2.png): Summary of the landscape conditions around the sampling points. Average percentage of surface occupied by each land cover class in the various buffer zones (250 m, 500 m, 1 km, 2 km radii) around the collection points. Error bars indicate the mean +/- one standard deviation considering all the sampling points.

Additional file 3 (add_file_3.pdf): Pictures representative of the main land cover classes in the Diébougou area. Pictures were taken in November 2018.

Additional figure 4 (add_fig_4.pdf): Model evaluation plots for the presence models. A1, A2, A3 are Precision-Recall curves for the presence models of respectively *An. funestus*, *An. Gambiae s.s.* and *An. coluzz*ii. The baseline PR curve for each model is represented by the horizontal dashed line. B1, B2, B3 are observed vs. predicted presence probabilities for each out-of-sample village. The y-axis represents the sum over the 8 sampling points / village / survey (4 points by village * 2 places (interior and exterior)).

Additional figure 5 (add_fig_5.pdf) : Model evaluation plots for the abundance models. A1, A2, A3 are violin plots of the distribution of the residuals for the abundance models of respectively *An. funestus*, *An. gambiae s.s.* and *An. coluzz*ii. Black dots indicate the median value. B1, B2, B3 are observed vs. predicted number of bites / village / entomological surveys. The y-axis represents the sum of bites over the 8 sampling points / village / survey (4 points by village * 2 places (interior and exterior)) on a logarithmic scale. The absence of a dot indicates that no vector was collected. MAE = Mean Absolute Error; n = number of observations

## Declarations

### List of abbreviations

IML: Interpretable machine learning; ML: machine learning; HLC: Human landing catch; VC : Vector control; RF: Random forest; cc: correlation coefficient; EO: Earth Observation; SPOT: Satellite Pour l’Observation de la Terre; SRTM: Shuttle Radar Topography Mission; DEM: Digital elevation model; LST: Land Surface Temperature; CCM: Cross-correlation map; PCR: Polymerase chain reaction; LVO-CV: leave-village-out cross-validation; PR-AUC: Precision- Recall Area Under Curve; MAE: Mean Absolute Error; VIP(s): Variable importance plot(s); PDP(s): Partial dependence plot(s); b\w: between; SD: standard deviation; NDVI: Normalize difference vegetation index

### Ethics approval and consent to participate

The protocol of this study was reviewed and approved by the Institutional Ethics Committee of the Institut de Recherche en Sciences de la Santé (IEC-IRSS) and registered as N°A06/2016/CEIRES. Mosquito collectors and supervisors gave their written informed consent. They received a vaccine against yellow fever as a prophylactic measure. Collectors were treated free of charge for malaria according to WHO recommendations.

### Consent for publication

Not applicable Availability of data and materials

The datasets used and/or analyzed during the current study are available from the corresponding author on reasonable request.

### Authors’ contributions

PT, CP and NM conceived and designed the study. DDS and PT collected and prepared the data. NM, KM and RKD surpervised fieldwork. PT analyzed the data, helped by AP and MM. PT and NM drafted the manuscript. KM, CP, DDS, AAK, RKD, MM and AP reviewed the manuscript. All authors read and approved the final manuscript.

## Supporting information

Additional figures (all)

## Acknowledgements

We thank populations of the villages for their kind support and collaboration. We also thank all the field and laboratory staff for their strong commitment to the REACT project. We thank all the anonymous persons that make data-science web forums alive by asking and answering technical questions; these forums were extensively used to elaborate this modeling work. Map data copyrighted OpenStreetMap contributors and available from https://www.openstreetmap.org. This work contains modified Copernicus Sentinel data (2018).

