## Additional figures (all) for "Data-driven and interpretable machine-learning modeling to explore the fine-scale environmental determinants of malaria vectors biting rates in rural Burkina Faso"

### Additional figure 1

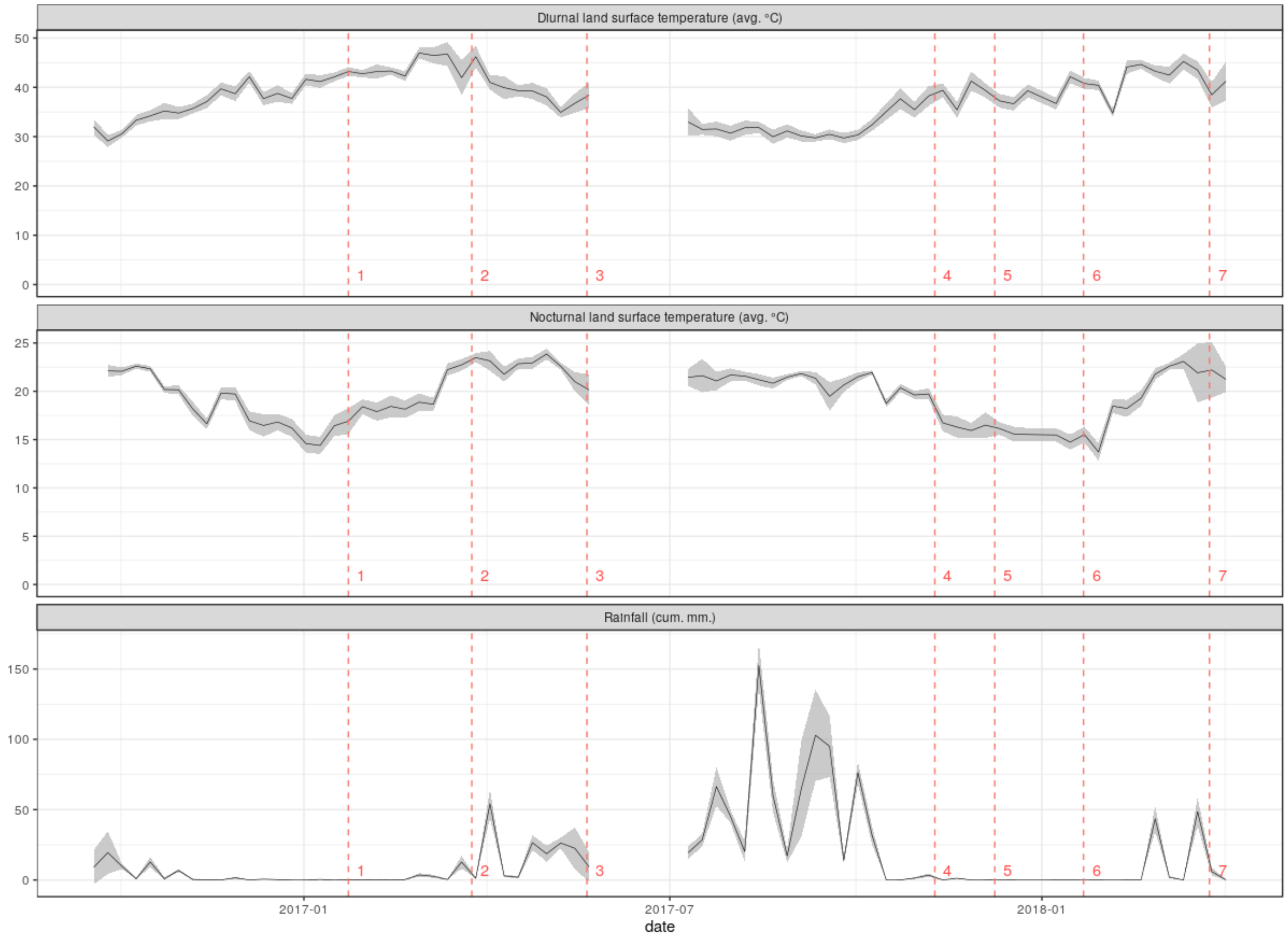


data sources : GPM (rainfall), MODIS (temperature)

### Additional figure 2

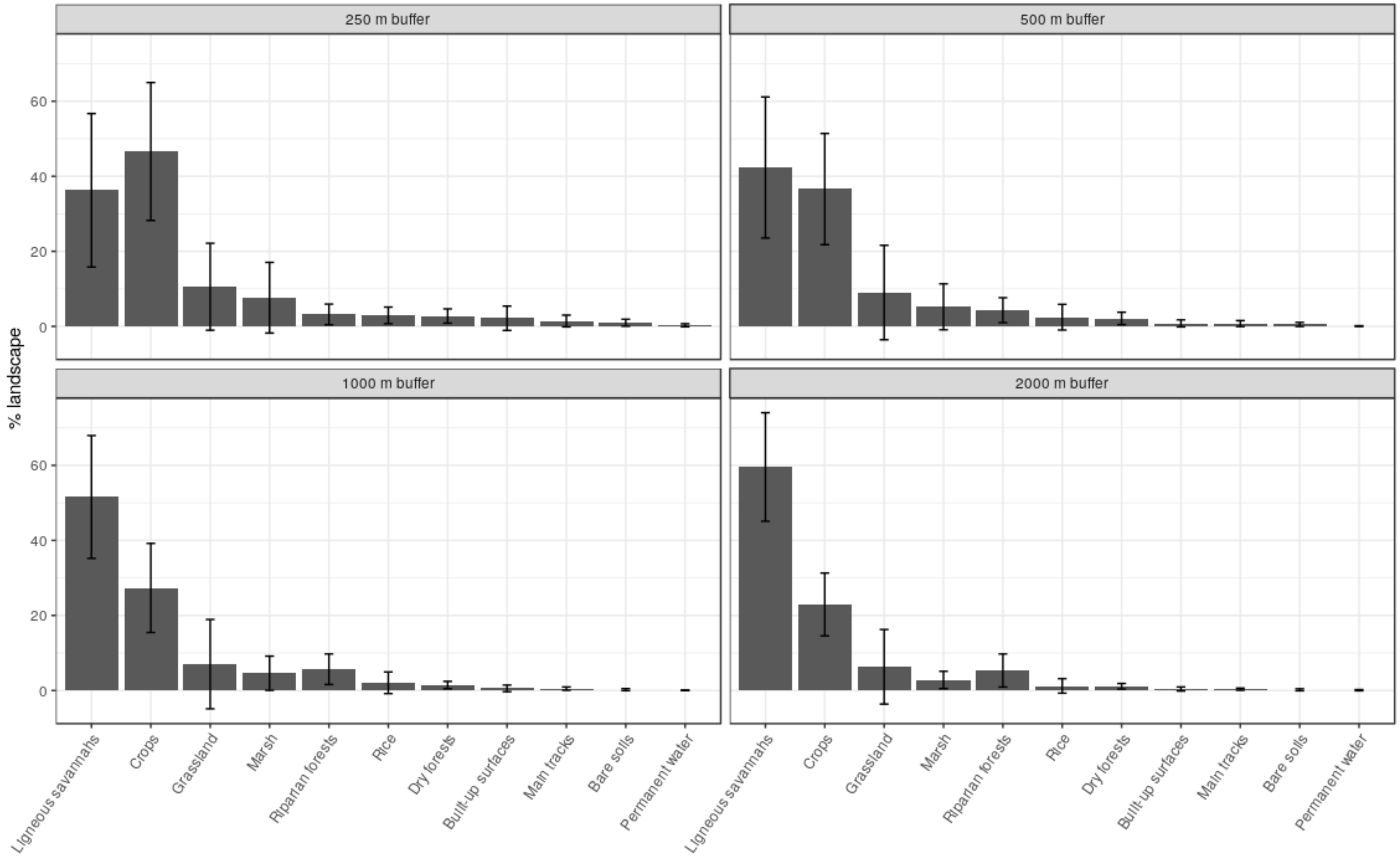


Data source : land cover map built from a supervised classification using satellite images

### Additional file 3

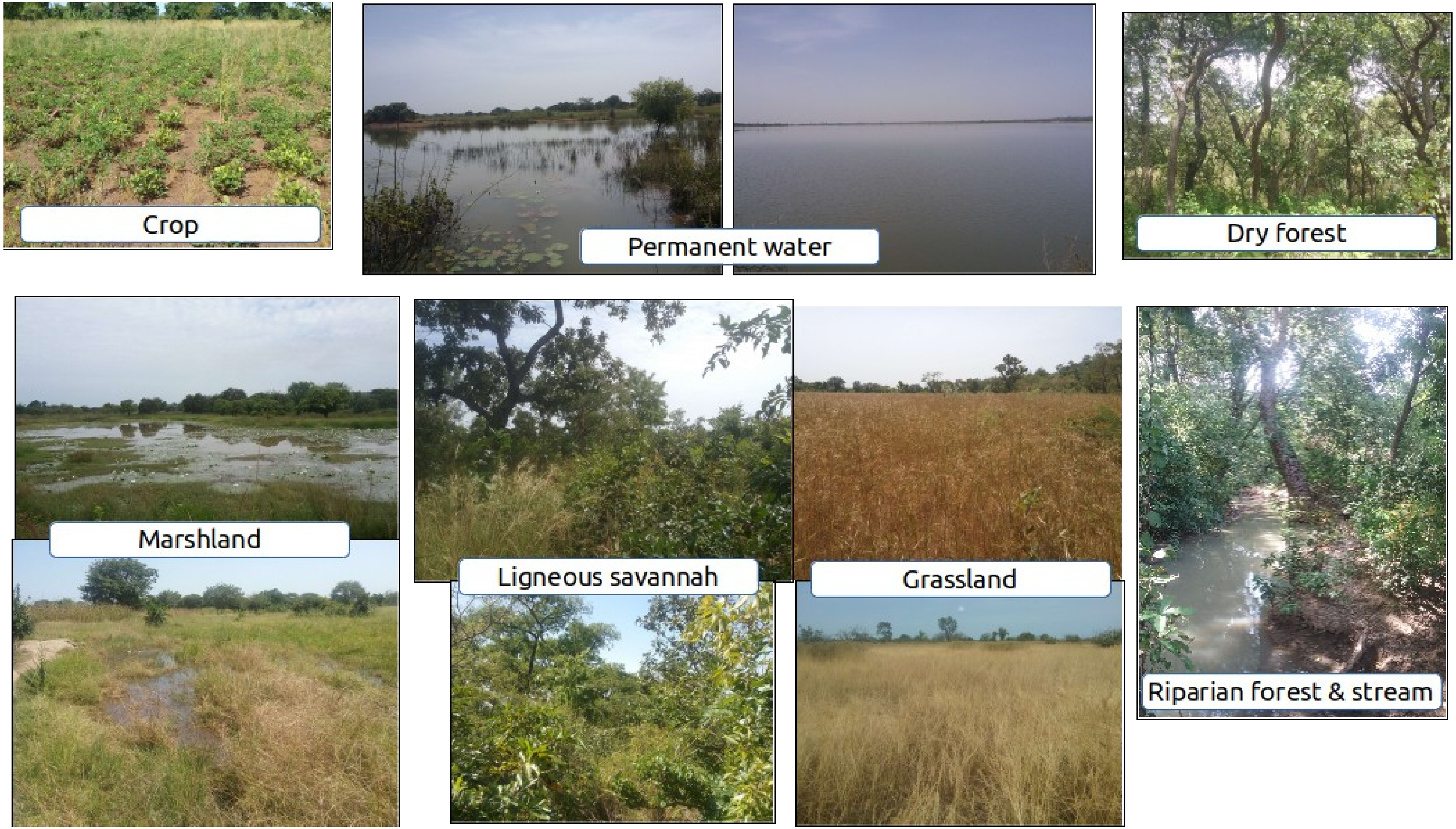

### Additional figure 4

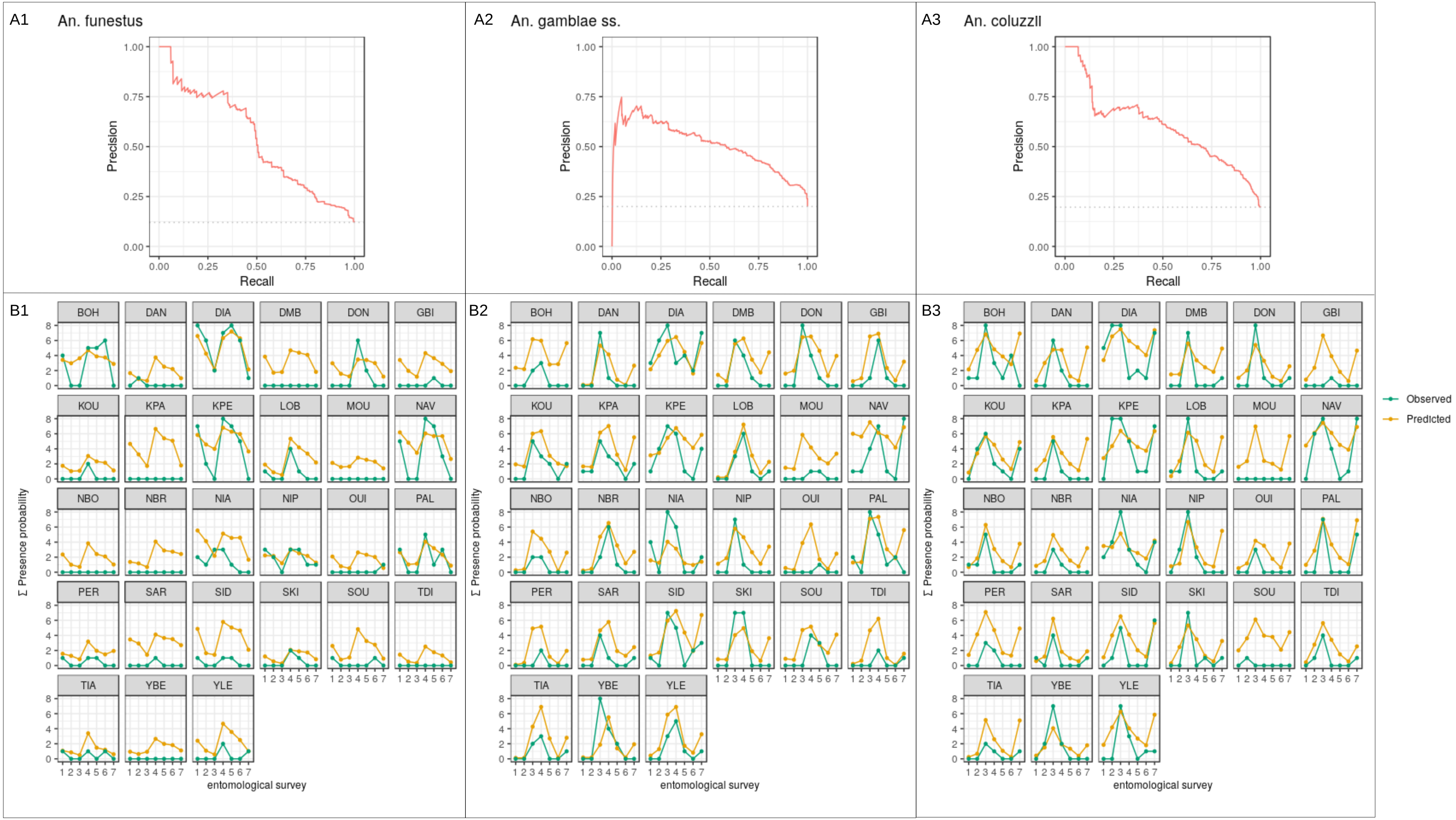

### Additional figure 5

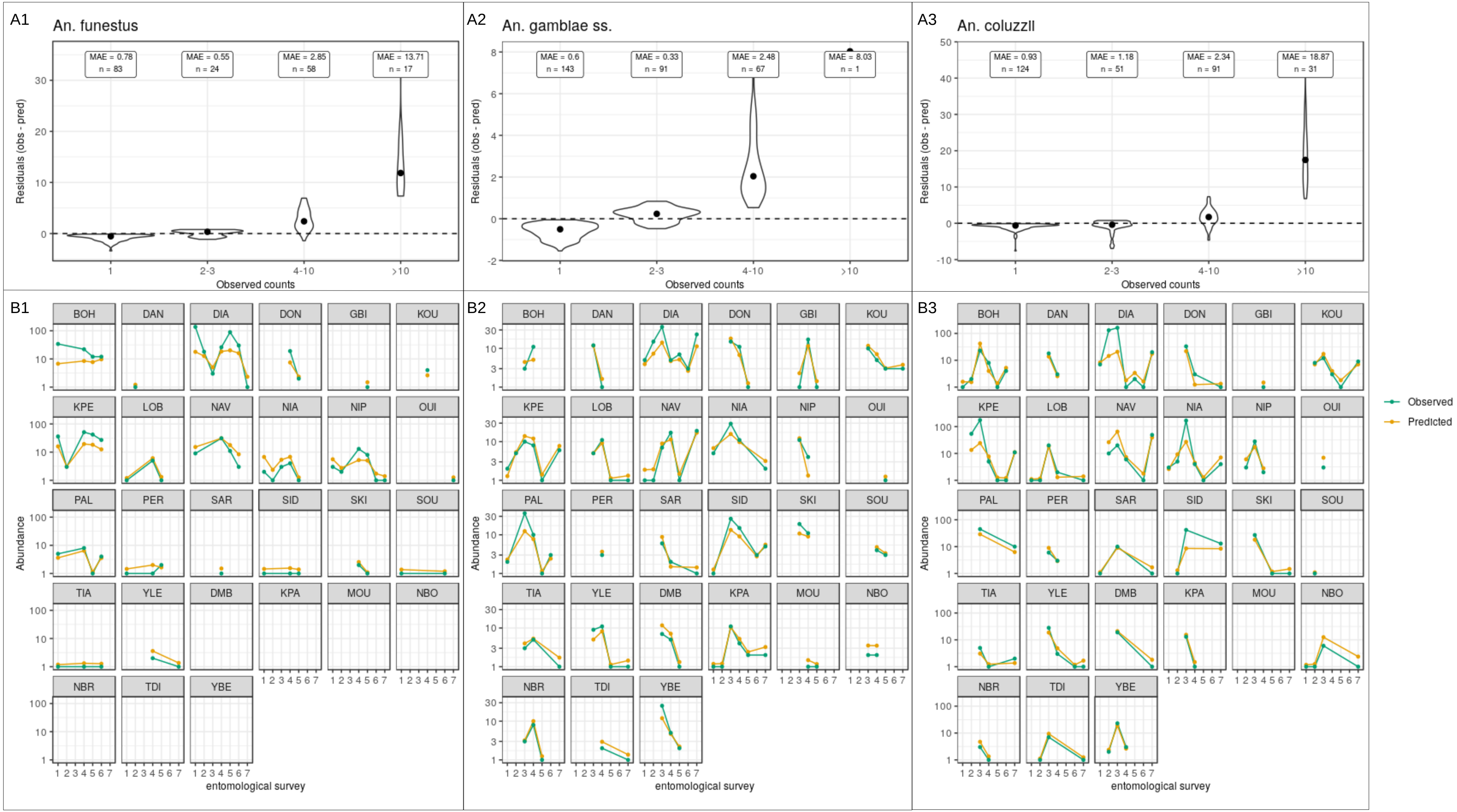
